# Genotype and methylation interact to reconfigure transcriptional regulation in colorectal cancer

**DOI:** 10.64898/2026.05.27.728350

**Authors:** Byounghun Kim, Hanbeen Kim, Min-Kyeong Kwon, Sridhar Hannenhalli, Sun Shim Choi

## Abstract

**Background:** Transcriptional regulation is shaped by both genomic variants and the environment. Yet, how the regulatory effects of genomic variants are reconfigured by dynamic epigenomic changes during tumorigenesis remains incompletely understood.

**Methods:** We investigated methylation context-dependent links between genotype and gene expression in colorectal cancer (CRC) using paired tumor and normal-adjacent tissue (NAT) from 80 patients, thereby controlling for germline genomic background. By integrating promoter-targeted bisulfite sequencing with RNA-seq, we systematically compared expression quantitative trait loci (eQTLs) and methylation quantitative trait loci (mQTLs). To capture regulatory complexity beyond simple mediation, we implemented a memo-eQTL framework that explicitly models genotype × DNA methylation (G×M) interactions.

**Results:** We observed extensive tissue specificity in both eQTL and mQTL landscapes; tumor-specific eGenes were significantly enriched for hallmark oncogenic pathways, including WNT and MAPK signaling. Standard mediation models explained only a minority of genotype–expression relationships, whereas our explicit interaction framework revealed widespread reconfiguration of methylation-dependent genetic effects in tumors. Memo-eQTL mapping (FDR < 0.05) identified 18 NAT and 73 tumor eGenes with significant G×M interactions, and results were consistent at a more permissive threshold (FDR < 0.2). We further developed a patient-level memo-eQTL score and found that interaction-based regulatory disruption in NAT, but not in tumor, significantly correlated with clinical stage (*P* = 0.035).

**Conclusions:** Genetic regulation in cancer is reorganized through context-dependent G×M interactions. Importantly, G×M signatures in NAT are specifically linked to disease progression, offering new insights into field cancerization and the clinical consequences of regulatory reprogramming in CRC.

## Background

Genomic variation contributes to disease susceptibility and progression not only through coding mutations but also through alterations in the non-coding regions of the genome ultimately affecting transcriptional regulation [1,2]. Consistently, a large proportion of disease-associated variants identified by genome-wide association studies reside in non-coding regulatory DNA, where they potentially modulate transcriptional and epigenetic processes [3,4]. In this context, expression quantitative trait locus (eQTL) and methylation quantitative trait locus (mQTL) analyses provide essential, quantitative links between inherited genomic variation and downstream regulatory phenotypes, enabling systematic study of how germline variants influence gene expression and DNA methylation across human tissues [5–7].

Although eQTLs and mQTLs capture statistically robust genotype–phenotype associations, these regulatory effects are not independent and frequently context-dependent; the same variant can have different regulatory consequences across tissues, cell types, developmental stages, or environmental exposures [8]. Recent single-cell and perturbation studies have shown that many genetic effects are cell-type or environment specific, motivating context-sensitive approaches for dissecting regulatory architectures [9–11].

During tumorigenesis, the epigenetic landscape is extensively remodeled, with widespread alterations in DNA methylation that can reconfigure regulatory relationships observed in normal tissue [12,13]. Colorectal cancer (CRC) is a particularly informative system for studying these dynamics due to pervasive epigenetic changes, such as CpG island hypermethylation (CIMP) [14,15]. Crucially, paired normal-adjacent tissue (NAT) and tumor samples share an identical germline background, enabling within-individual comparisons that minimize confounding by inter-individual genetic variation.

While NAT has traditionally been used as a baseline ‘healthy’ control, it is becoming clearer that they are distinct from healthy tissue, and our previous work demonstrated that NAT-derived transcriptomes are significantly more informative than tumor-derived transcriptomes in predicting CRC recurrence [16]. We have previously shown that NATs exhibit a greater number of differentially expressed genes (DEGs) between recurrent and non-recurrent cases compared to tumors. Furthermore, prognostic models built on NAT-derived data outperformed tumor-based models in independent validation cohorts, such as TCGA-COAD, and revealed superior prognostic capability in tumor-infiltrating immune cell compositions [16].

In the present study, we aim to elucidate how inherited variations contribute to cancer phenotypes by (i) comparing regulatory effects across donor-matched tissue states controlling for germline background, and (ii) utilizing analytical frameworks that explicitly capture context-dependent regulation. Focusing on promoter-proximal regulation, we mapped promoter-proximal eQTLs and mQTLs in both NAT and tumor tissue, subsequently applying a memo-eQTL mapping framework to test for significant genotype × DNA methylation (G×M) interactions [17]. Leveraging paired multi-omic samples from the same 80 CRC patients used in our previous transcriptomic study [16], we quantified the reconfiguration of promoter-proximal genetic regulation during tumorigenesis. Finally, we examined whether interaction-based regulatory disruption, specifically in the NAT or tumor environments, relates to clinical heterogeneity, providing a deeper mechanistic link between germline variation and disease progression.

## Methods

### Study cohort and data overview

We analyzed paired NAT and tumor samples from 80 Korean CRC patients recruited at Samsung Medical Center, as described in our previous work [16]. From each sample we generated promoter-targeted bisulfite sequencing data to quantify CpG methylation within gene promoters [18] and matched RNA-seq data to quantify gene expression [16]. These paired multi-omics data including promoter methylation, expression, and bisulfite-derived genotypes enabled integrated, within-individual analyses of genotype, DNA methylation, and transcription across NAT and tumor tissues.

### Bisulfite sequencing data preprocessing

Promoter-targeted bisulfite sequencing reads were preprocessed for both methylation quantification and SNP genotyping. Raw reads were first quality-checked with FastQC (v0.11.9) [19], and adapters and low-quality bases were removed using Trim Galore (v0.6.7) [20]. Cleaned reads were aligned to the GRCh38 reference genome using Bismark (v0.22.1), a bisulfite-aware aligner that accounts for C to T conversions introduced by bisulfite treatment [21]. PCR duplicates were removed using Bismark’s deduplication routine. Base-quality recalibration and SNP-aware processing were performed with bis-SNP to improve the accuracy of variant calls from bisulfite data [22].

Downstream SNP calling referenced the Korea 1K genome dataset to represent population-level variation in Koreans [23]. SNVs were retained if the genotype was concordant between NAT and tumor samples (i.e., 0/0, 0/1, or 1/1 in both) within at least one patient, thereby restricting analyses to putative germline variants. Further filtering required that SNVs be present in the Korea 1K reference set. We then restricted SNVs to promoter regions, defined as −1,500 to +200 bp relative to the transcription start site (TSS) of each gene [24]. Rare variants were excluded using a minor-allele frequency (MAF) threshold of 1%. After all filtering steps, 17,013 promoter SNVs were retained for QTL mapping.

### RNA sequencing data preprocessing and gene expression quantification

RNA sequencing data were preprocessed for gene expression quantification. An index of the GRCh38 reference genome was generated using the STAR aligner (v2.7.10) and used consistently for all subsequent RNA-seq read alignments [25]. Quality control of raw RNA-seq reads was performed using FastQC (v0.11.9), followed by removal of adapter sequences and low-quality bases when necessary [19]. The cleaned reads were aligned to the reference genome using STAR with splice junction-aware alignment to accurately capture exon–intron structures. After alignment, gene-level read counts were obtained using HTSeq-count based on GENCODE v40 gene annotation [4,26]. Gene expression levels were quantified as FPKM (Fragments Per Kilobase of transcript per Million mapped reads) values calculated from the raw read counts to enable comparison across samples [27]. To ensure robustness in downstream QTL analyses, only genes detected in at least 30% of all samples and annotated as protein-coding with defined transcription start sites (TSS) were retained. After filtering, a total of 15,667 genes were included in downstream analyses.

### QTL mapping

All QTL analyses were conducted separately for NAT and tumor samples. For each gene, the promoter was defined as −1,500 to +200 bp relative to TSS, and SNPs within this window were considered candidate promoter-proximal QTL variants. In eQTL analysis, gene expression was treated as a continuous outcome. We tested SNP–phenotype associations using linear regression models of the form:

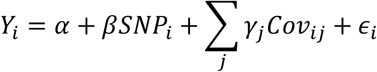

where *Y*_*i*_ is the phenotype (gene expression or CpG methylation) for sample *i*, *SNP*_*i*_ denotes the genotype dosage for *sample*_*i*_, and *β* represents the effect size of the SNP, and *Cov*_*ij*_ represents the j^th^ covariate for *sample*_*i*_with *γ*_*j*_ being the corresponding coefficient. Covariates included age, sex, and anatomical location of the sample within the colon (eight categories: ascending colon, transverse colon, descending colon, sigmoid colon, rectum, rectosigmoid junction, hepatic flexure, and splenic flexure), along with latent factors estimated with PEER (Probabilistic Estimation of Expression Residuals) to account for unobserved confounding [28]. Note that the covariates are shared by both NAT and tumor tissues obtained from the same donor. The error term *∈*_*i*_ captures residual variation unexplained by the model. We assessed multicollinearity among covariates and found no evidence of problematic collinearity. Promoter-proximal eQTL mapping was performed with FastQTL (v2.0) and promoter-proximal mQTL mapping with tensorQTL (v1.0.10), a computationally efficient implementation of permutation-based QTL mapping [29,30]. Nominal association tests were computed for all SNP–phenotype pairs, followed by permutation testing to obtain gene- (or promoter-) level adjusted *P*-values [31]. SNP–phenotype pairs with permutation-based adjusted *P* < 0.05 were considered significant promoter-proximal eQTLs or mQTLs.

### Variance-matched permutation test

To further account for potential variance-related bias in pathway enrichment, we implemented a variance-matched permutation framework. All genes in the expression matrix were stratified into deciles based on their expression variance in tumor tissue. We then assessed the enrichment of WNT and MAPK signaling pathways within the tumor eGene set using a Fisher’s exact test. To generate a null distribution, we created 10,000 random gene sets by sampling from the background gene pool while strictly maintaining the same variance decile profile as the observed tumor eGenes. A permutation *P*-value was calculated as the proportion of these 10,000 random trials that yielded an enrichment *P*-value equal to or more significant than the observed value for each pathway. This approach explicitly tests whether the observed enrichment is a unique feature of the eQTL landscape or a generic consequence of sampling high-variance genes.

### Memo-eQTL identification

We implemented memo-eQTL mapping by fitting three nested linear regression models for each ‘SNP–CpG–gene’ triplet, following the framework described previously [17]. Genotypes were encoded under an additive model (major allele homozygote = 0, heterozygote = 1, minor allele homozygote = 2).

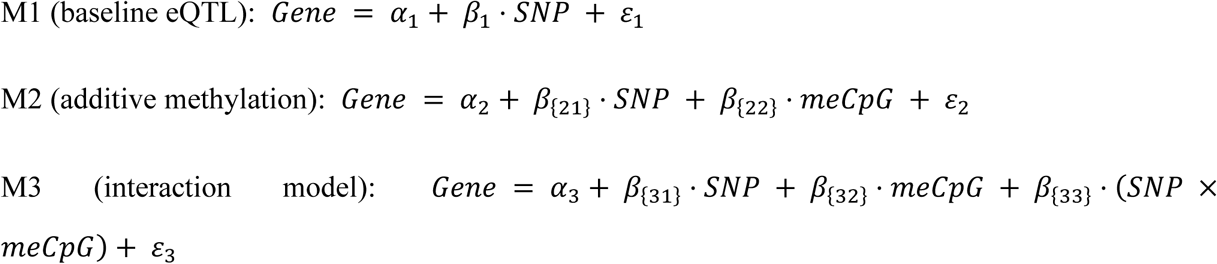

All models included the same covariates used in the eQTL analyses (age, sex, tumor location, and PEER factors), and analyses were performed independently for NAT and tumor tissues. Methylation at each CpG site was analyzed individually, and measurements with sequencing coverage < 10 reads in a given sample were set to missing. For each triplet-level regression, samples with missing genotype, methylation, expression, or covariate values were excluded; analyses were performed only when at least 50 samples remained after filtering to ensure sufficient statistical power. To identify methylation-modulated genetic effects, we compared nested models using likelihood ratio tests (LRTs). Specifically, M3 was compared to M2 (df = 1) to assess the contribution of the SNP × methylation interaction term, and to M1 (df = 2) to evaluate the combined contribution of methylation and interaction terms. *P*-values derived from these tests were corrected for multiple testing using the Benjamini–Hochberg false discovery rate (FDR) procedure across all tested triplets within each tissue. Triplets were defined as significant memo-eQTLs if both comparisons (M3 vs M2 and M3 vs M1) were significant after FDR correction (FDR < 0.05). In addition, a more permissive threshold (FDR < 0.2) was applied to identify a broader set of candidate memo-eQTLs, which are reported in the Supplementary Table S4.

### Quantification of memo-eQTL interaction effects by triplet

For each significant memo-eQTL (SNP–meCpG–gene triplet), we computed an interaction component (IC) to quantify the per-patient magnitude and direction of the SNP × methylation interaction. The IC for triplet t in patient i was defined as:

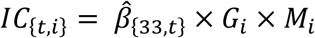

where *β̂*_{33,*t*}_ is the estimated SNP × meCpG interaction coefficient from model M3 for triplet *t*, *Gi* is the genotype dosage for patient *i* (0, 1, 2), and *Mi* is the methylation measurement for the corresponding meCpG in patient *i*. Prior to IC calculation, CpG methylation values were z-score normalized across samples (mean = 0, SD = 1). IC values were set to missing for any patient–triplet pair in which the interaction coefficient could not be estimated or genotype/methylation data were missing.

To compare interaction effects between NAT and tumor, we constructed tissue-specific IC matrices using the same set of triplets (those that were significant in either NAT or in tumor). Each matrix had triplets as rows and patients as columns (rows = triplets; columns = patients). For visualization of interaction strength, heatmaps were generated using row-wise z-score–normalized interaction components, such that for each triplet, IC values were standardized across patients and tissues.

### Calculating patient-level memo-eQTL interaction scores and clinical association analysis

For each significant memo-eQTL triplet in either NAT or in tumor, we computed an IC per patient as described above. Using these IC values, we defined an individual memo-eQTL magnitude score to summarize the overall burden of genotype × methylation interaction effects carried by each patient in a given tissue.

For patient *i* and tissue *k*, the memo-eQTL magnitude score was calculated as the mean absolute IC across all memo-eQTL triplets measured in that tissue:

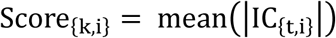

The mean was calculated using non-missing IC values for each patient, without applying additional weighting across triplets.

To quantify changes in interaction strength between NAT and tumor tissue, differences in magnitude scores between tissues were computed using the same scoring framework. For memo-eQTL sets defined in normal tissue (NAT memo-eQTL) and tumor tissue (tumor memo-eQTL), the following difference score was calculated:

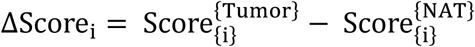

The ΔScore_i_ calculated using memo-eQTL triplets identified in NAT reflects the extent to which interaction patterns defined in normal tissue are altered in tumor tissue.

To compare ΔScore_i_ magnitudes by tumor progression, patients were dichotomized into two groups: stages I–II (group 1) and stage III (group 2). We then fitted univariate logistic regression with Tumor-Node-Metastasis (TNM) stage (group 2 vs. group 1) as the outcome and the individual ΔScore_i_ magnitude as the predictor. Results are reported as odds ratios with 95% confidence intervals, and statistical significance was assessed using the Wald test.

### Motif-based analysis of memo-eQTL CpGs

To characterize transcription factor (TF) involvement in methylation-modulated regulatory interactions, we annotated TF binding sites (TFBS) across the genome using position weight matrices (PWMs) obtained from the JASPAR core database [32]. Genome-wide motif scanning was performed on the GRCh38 reference genome using a PWM-scan tool [33], and candidate TFBS were identified using a default affinity score threshold. CpG sites located within ±100 bp of predicted TFBS were defined as TF-associated CpGs. Motif identifiers (MA_ID) were mapped to TF symbols using a curated JASPAR motif-to-gene annotation table.

## Results

### Study design and cohort overview

The primary goal of this study was to determine how G×M interactions shaping gene expression are reconfigured between normal and tumor states. To address this, we integrated promoter DNA methylation, gene expression, and promoter-centered genotypes (bisulfite-derived SNPs) from paired NAT and tumor samples of 80 Korean CRC patients (Figure 1A; see Methods).

**Figure 1.**
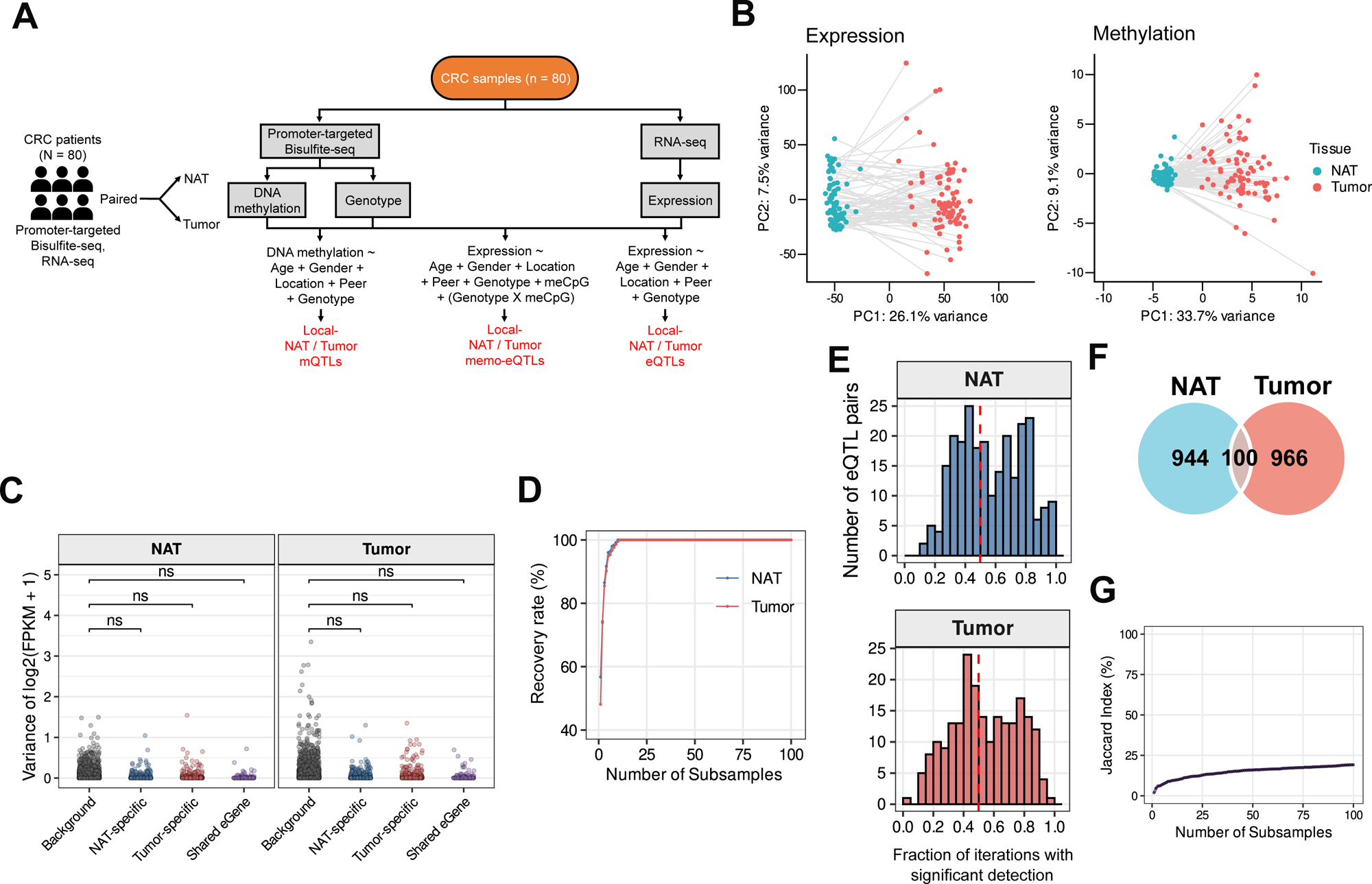
Study design and tissue-specific eQTL landscapes in colorectal cancer. (**A**) Schematic illustration of the study design and analytical framework. Paired normal adjacent tissue (NAT) and tumor samples from 80 colorectal cancer patients were profiled using promoter-targeted bisulfite sequencing and RNA sequencing to obtain genotype, DNA methylation, and gene expression data. These data were used to perform promoter-proximal mQTL, eQTL, and methylation-modulated eQTL (memo-eQTL) analyses. (**B**) Principal component analysis (PCA) of RNA-seq–based gene expression data (left) and promoter-targeted DNA methylation data (right). Each point represents an individual sample, with paired NAT and tumor samples from the same patient connected by grey lines. Colors indicate tissue type. (**C**) Expression variance distributions of eGenes and background genes. For each tissue (NAT, left; Tumor, right), the variance of log_2_(FPKM + 1) was calculated per gene across 80 samples. eGenes were stratified into NAT-specific (blue), tumor-specific (red), and shared (purple) categories and compared against background genes (non-eGenes) using two-sided Wilcoxon rank-sum tests. ns, not significant. (**D**) Saturation analysis of eQTL discovery. The ground-truth eQTL set was defined from the full cohort (N = 80). In each of 100 bootstrap iterations, 80% of individuals (n = 64) were randomly subsampled, and significant eQTLs were cumulatively merged. The recovery rate represents the proportion of ground-truth eQTLs recovered by the cumulative union at each iteration. Blue, NAT; red, tumor. (**E**) Per-eQTL reproducibility across bootstrap iterations. For each significant eQTL pair identified in the full cohort (N = 80), the fraction of 100 bootstrap iterations (80% subsampling, N = 64) in which it was significantly detected was calculated. Histograms show the distribution of per-eQTL reproducibility fractions for NAT (left) and Tumor (right). Red dashed lines indicate the 0.5 reproducibility threshold. (**F**) Overlap of promoter-proximal eQTLs (SNP–gene pairs) identified in NAT and tumor tissue. (**G**) Persistence of tissue-specific eQTL landscapes across bootstrap iterations. At each iteration, cumulative eQTL unions were constructed independently for NAT and tumor tissues, and the Jaccard index between the two cumulative SNP–gene pair unions was calculated.

To assess the global molecular landscape of our cohort, we performed principal component analysis (PCA) on both gene expression and promoter methylation profiles. In both assays, PC1, explaining 26.1% of expression variance and 33.7% of methylation variance, clearly separates samples by tissue state (Figure 1B). As expected, NAT samples form a tight, homogeneous cluster, while tumor samples exhibit substantially greater dispersion, reflecting the characteristic transcriptional and epigenetic heterogeneity of tumors. The tumor-associated regulatory shifts captured by PC1 substantially exceed baseline inter-individual variation, underscoring the profound impact of the tumor environment on molecular phenotypes. These clear molecular distinctions provide a strong rationale for treating NAT and tumor as separate regulatory environments in all subsequent tissue-specific QTL and G×M interaction analyses.

As shown in Figure 1A, we used this promoter-centric multi-omics resource to perform three complementary cis-regulatory mapping analyses within NAT and tumor tissue contexts: cis-mQTL mapping (methylation ∼ genotype + covariates) to assess genetic effects on DNA methylation; cis-eQTL mapping (expression ∼ genotype + covariates) to assess genetic effects on gene expression; and methylation-modulated eQTL mapping (expression ∼ genotype + methylation + G×M + covariates; memo-eQTLs) to identify genetic effects specifically modulated by local DNA methylation levels [17]. Complete lists of eQTLs and mQTLs identified in NAT and tumor tissues are provided in Supplementary Table S1 and S2.

To ensure the reliability of our eQTL calls, we performed two complementary robustness analyses. First, to exclude the possibility that eQTL detection was biased toward high-variance genes, we compared expression variance distributions between eGenes and background genes (genes not associated with any eQTL) using a two-sided Wilcoxon rank sum test. eGenes were stratified into three subclasses, i.e., NAT-specific, tumor-specific, and shared, and no significant difference in expression variance was observed between any eGene subclass and background genes in either tissue (Figure 1C). Second, to assess whether eQTL detection was robust given the modest sample size (n = 80), we performed a bootstrap recovery analysis. Across 100 iterations of independent subsampling, randomly selecting 80% of individuals (n = 64) per iteration, significant eQTL lists from each subsample were sequentially merged to form a cumulative union, and the proportion of the full eQTL list recovered was tracked across iterations. The cumulative recovery rate reached approximately 95% within 15 bootstrap iterations in both NAT and tumor tissues and plateaued near 100% by 25 iterations (Figure 1D). The near-identical convergence trajectories in NAT and tumor indicate that eQTL detection stability is comparable across tissue states, confirming that our eQTL calls are not sensitive to the specific composition of the patient cohort.

To further assess stability at the level of individual eQTL pairs, we computed the fraction of 100 bootstrap iterations in which each eQTL was significantly detected (Figure 1E). In NAT, 57.1% of eQTLs (144 of 252) were reproduced in more than half of iterations, and 18.3% (46 of 252) were detected in over 80% of iterations (Figure 1E, top panel). In tumor, 52.3% (112 of 214) exceeded the 50% threshold, and 14.5% (31 of 214) exceeded 80% (Figure 1E, bottom panel). The nearly identical distribution patterns between NAT and tumor confirm comparable per-eQTL detection stability across tissue states.

Having established the robustness of our eQTL calls, we examined the degree of tissue-specificity across the full cohort. Comparison of the NAT and tumor regulatory maps revealed a striking degree of tissue-specificity: despite the identical germline background of the paired samples, only a small fraction of eQTLs (n = 100) were shared between tissues, while the vast majority were identified exclusively in one tissue state, 944 in NAT and 966 in tumor (Figure 1F; Supplementary Table S1). To further verify that the limited inter-tissue eQTL overlap is not an artifact of finite sample size, we performed a cumulative inter-tissue overlap analysis based on the bootstrap subsampling described above. For each iteration, cumulative unions of eQTL pairs were independently assembled for NAT and tumor tissues, progressively expanding the pool of identifiable eQTL candidates. The Jaccard index between the cumulative NAT and tumor eQTL unions remained consistently low (∼20 %) and stable across all 100 iterations (Figure 1G), confirming that the limited overlap between NAT and tumor eQTLs reflects genuine tissue-specific regulatory divergence rather than a sampling artifact.

### Tissue-specificity of promoter-proximal eQTLs

We next compared the estimated eQTL effect sizes (regression coefficients, β) to quantify the magnitude of regulatory shifts (Figure 2A–C). NAT-specific eQTLs exhibited large absolute effect sizes in NAT that were markedly attenuated in tumor tissue (Figure 2A), whereas tumor-specific eQTLs showed the converse pattern (Figure 2B). Shared eQTLs largely maintained concordant effect directions across both tissues (Figure 2C). The regulatory effects of NAT-specific and tumor-specific eQTLs were consistently attenuated or lost in the other tissue environment, confirming that these variants operate under strict context-dependent logic. Examples of individual loci further highlight these patterns: NAT-specific eQTL (e.g., *SRI*) exhibiting a clear genotype-dependent expression gradient in normal tissue that disappears in the tumor (Figure 2D). Tumor-specific (e.g., *BEST1*) exhibiting a ‘gain-of-function’ regulatory effect where the SNP only influences expression in the malignant state (Figure 2E). Shared eQTLs (e.g., *PSMD13*) presenting in both tissues but showing altered slopes or expression baselines, suggesting that even shared effects undergo partial reconfiguration (Figure 2F).

**Figure 2.**
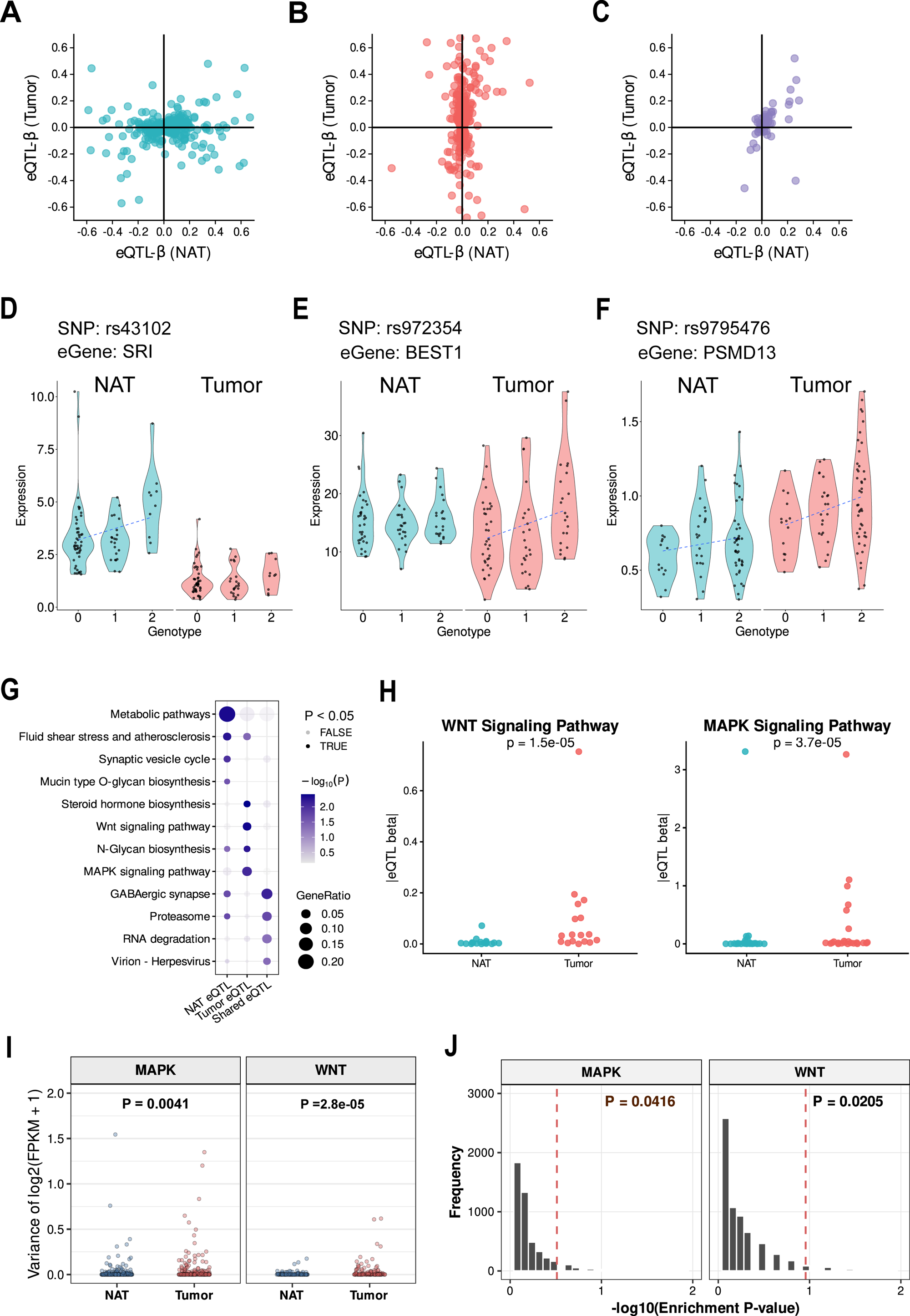
Tissue-specific promoter-proximal eQTLs, pathway enrichment, and variance-matched validation. (**A–C**) Comparison of eQTL effect sizes (regression coefficients, β) between NAT and tumor tissues. Scatter plots show eQTL regression coefficients estimated in NAT (x-axis) and tumor (y-axis) for NAT-specific eQTLs (**A**), tumor-specific eQTLs (**B**), and shared eQTLs (**C**). Each point represents an individual eQTL. (**D–F**) Representative examples of NAT-specific (**D**), tumor-specific (**E**), and shared (**F**) eQTLs. Violin plots show genotype-dependent gene expression levels in NAT and tumor tissues for each example locus. Points represent individual samples. (**G**) Pathway enrichment analysis of eGenes associated with NAT-specific, tumor-specific, and shared eQTLs. Dot size indicates gene ratio, and color represents −log₁₀(adjusted P-value). (**H**) Comparison of eQTL effect sizes for WNT and MAPK signaling pathway genes between NAT and tumor tissues. Each point represents a pathway gene, and absolute eQTL effect sizes (|β|) in NAT and tumor are shown as separate groups. (**I**) Comparison of expression variance for MAPK and WNT signaling pathway genes between NAT and tumor tissues. The variance of log₂(FPKM + 1) was computed per gene across 80 samples. P-values were determined by two-sided Wilcoxon rank-sum tests. (**J**) Variance-matched permutation test for pathway enrichment. 10,000 random gene sets were generated by sampling from background genes while preserving the variance decile distribution of the observed tumor eGenes. Histograms represent the null distribution of −log₁₀(Fisher’s exact test P-values). Red dashed lines indicate the observed enrichment P-values. Permutation P-values are shown.

To assess the biological impact of this reconfiguration, we performed pathway enrichment analysis on eGenes (genes harboring eQTLs) associated with each tissue category (Figure 2G; Supplementary Table S3). NAT-specific eGenes were primarily involved in basal metabolic processes. In contrast, tumor-specific eGenes were significantly enriched for hallmark oncogenic drivers, most notably WNT and MAPK signaling. Consistently, WNT and MAPK pathway genes were observed to have significantly larger effect size in tumor compared to NAT (Figure 2H).

We next asked whether the observed enrichment of tumor eGenes in MAPK and WNT signaling pathways could be attributable to higher expression variability of these pathway genes in tumor tissue. Indeed, both MAPK and WNT pathway genes exhibited significantly higher expression variance in tumor compared to NAT (Figure 2I; MAPK: *P* = 4.1×10⁻^3^, WNT: *P* = 2.8×10⁻⁵), confirming that the concern raised above is legitimate in this dataset. To directly test whether this variance difference drove the observed pathway enrichment, we applied a variance-matched permutation framework in which 10,000 random gene sets were generated with the same variance decile composition as the observed tumor eGene set (see Materials and Methods). The enrichment of both WNT and MAPK pathways in tumor eGenes remained significant relative to this variance-matched null distribution (Figure 2J; MAPK: permutation *P* = 0.0416, WNT: permutation *P* = 0.0205), demonstrating that the pathway enrichment reflects genuine genotype-driven regulatory reconfiguration in tumor tissue rather than a statistical artifact of elevated expression variance.

Collectively, these results demonstrate that tumor-specific eQTLs are not random disruptions but are biologically targeted toward pathways relevant to the disease.

### Reconfiguration of mQTLs and the limits of simple DNA methylation mediation model

To determine if the tissue-specific shifts observed in eQTLs are mirrored in the epigenetic landscape, we independently mapped promoter-proximal mQTLs in NAT and tumor tissues (Figure 3A). Similar to our eQTL findings, the genetic cis regulation of DNA methylation is extensively reconfigured during tumorigenesis. While 63 mQTLs were shared between tissue states, a substantial majority were tissue-specific, with 643 detected exclusively in NAT and 409 identified only in tumor tissue (Supplementary Table S2).

**Figure 3.**
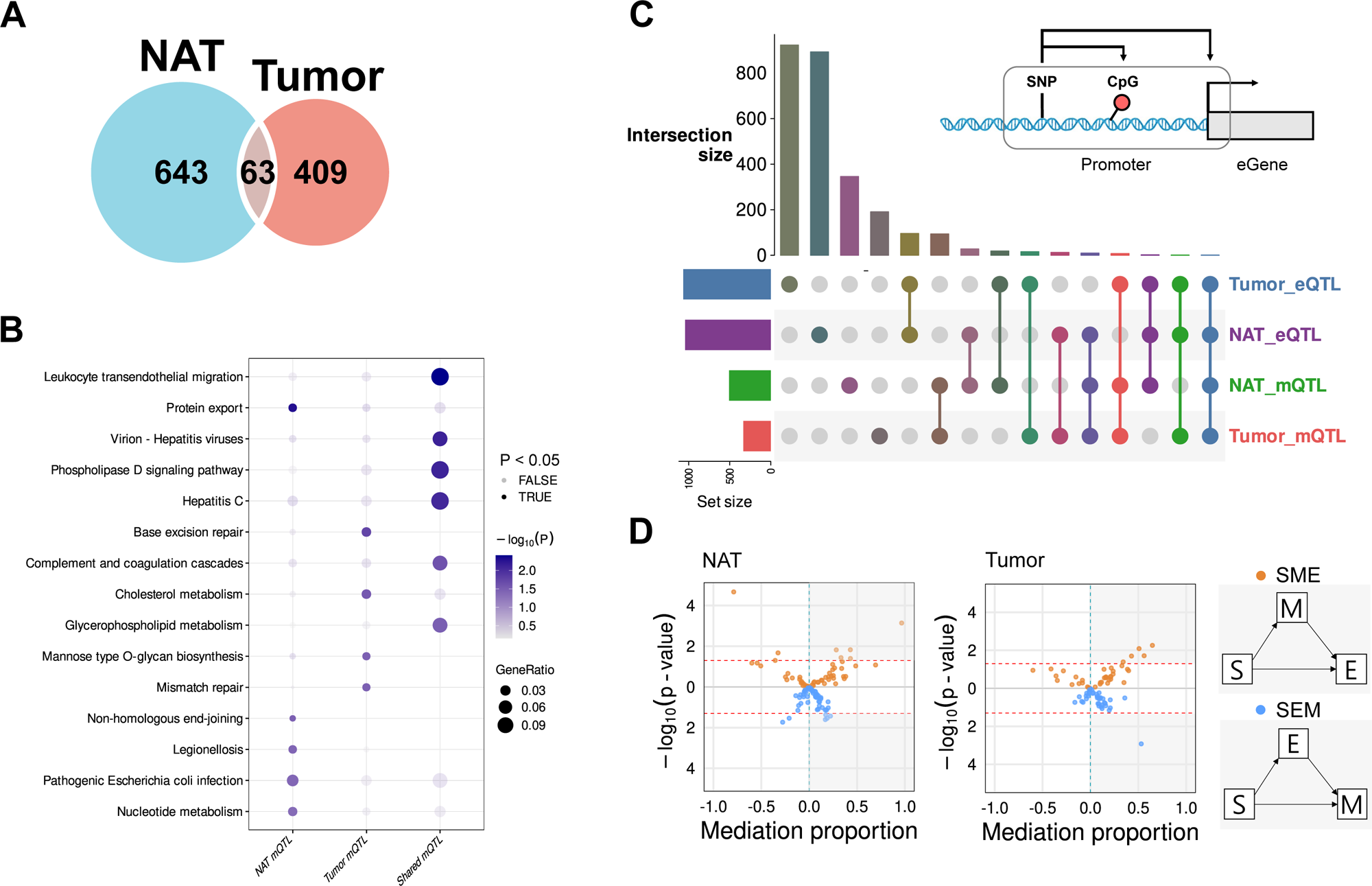
Tissue-specific promoter-proximal mQTLs and SNP–CpG–gene triplet configurations. **(A**) Overlap of significant promoter-proximal mQTLs (SNP–CpG pairs) identified in NAT and tumor tissues. (**B**) Pathway enrichment analysis of genes associated with NAT-specific, tumor-specific, and shared mQTLs. Dot size indicates gene ratio, and color represents −log10(adjusted *P*-value). (**C**) UpSet plot showing the overlap patterns of SNP–CpG–gene triplets across NAT eQTLs, tumor eQTLs, NAT mQTLs, and tumor mQTLs. Bars indicate the number of triplets for each combination. The nested schematic (top right) illustrates the structure of SNP–CpG–gene triplets used to examine the joint presence of eQTL and mQTL signals at promoter regions. (**D**) Mediation analysis of SNP–CpG–gene triplets in NAT (left) and tumor (right) tissues. Scatter plots show mediation proportion on the x-axis and −log₁₀(*P*-value) on the y-axis. Two directional mediation models were tested: SME (SNP → methylation → expression) and SEM (SNP → expression → methylation). Orange and blue points represent the SME and SEM models, respectively. Horizontal dashed lines indicate nominal (*P* < 0.05, inner) and FDR-corrected (outer) significance thresholds.

Pathway enrichment analysis of mQTL-associated genes revealed distinct functional landscapes across tissue categories (Figure 3B; Supplementary Table S4). NAT-specific mQTL genes were enriched for DNA metabolism and repair processes, including nucleotide metabolism and non-homologous end-joining, suggesting that genetic variants preferentially regulate the methylation of genomic maintenance genes in normal colonic tissue. In contrast, tumor-specific mQTL genes were significantly enriched for cancer-relevant DNA repair pathways, most notably base excision repair and mismatch repair, both of which are critically implicated in CRC pathogenesis. Shared mQTL genes were enriched for immune-related processes including leukocyte transendothelial migration and complement and coagulation cascades, reflecting constitutive regulatory relationships that persist across tissue states.

Notably, the functional categories enriched among mQTL-associated genes were largely distinct from those enriched among eQTL-associated genes, suggesting that genetic variants controlling promoter methylation and those controlling gene expression operate predominantly on different target genes. This functional divergence implies that eQTL SNPs and mQTL SNPs are largely independent in their regulatory consequences, providing a first indication that simple linear mediation, in which a single SNP drives both methylation and expression changes on the same gene, may not be the predominant regulatory architecture in this system.

To formally test this, we defined ‘SNP–CpG–gene’ triplets as instances where a SNP showed statistically significant associations with both gene expression (eQTL) and CpG methylation (mQTL), based on independent linear regression analyses (expression ∼ SNP and methylation ∼ SNP, respectively) (Figure 3C, nested panel). Triplets were constructed when the SNP had significant effect size estimates in both analyses. Analysis of these triplets using an UpSet plot revealed complex and largely non-overlapping regulatory patterns across tissue categories (Figure 3C). Triplets where all four regulatory signals (NAT eQTL, tumor eQTL, NAT mQTL, and tumor mQTL) co-occurred were rare, further underscoring that the coordinated genetic control of expression and methylation is highly sensitive to the cellular environment.

For triplets exhibiting both eQTL and mQTL signals, we applied mediation analysis to formally test whether the genotype–expression relationship is primarily driven by linear causal pathways (Figure 3D) [34]. We tested two directional models: (i) SME (SNP → Methylation → Expression), where methylation mediates genetic effects on expression, and (ii) SEM (SNP → Expression → Methylation), where expression mediates genetic effects on methylation. In both NAT and tumor tissues, only a small fraction of triplets reached statistical significance for either mediation model (Figure 3D; horizontal lines indicate nominal and FDR thresholds). The widespread lack of significant mediation, despite the presence of both significant eQTL and mQTL effects in these triplets, is consistent with the functional divergence observed between eGenes and mGenes, and indicates that simple linear models often fail to capture the true regulatory relationship between genotype, methylation, and expression. These observations motivated us to adopt an interaction framework in which promoter methylation modulates genotype effects in a context-sensitive manner, rather than acting as a fixed intermediate.

### Methylation-modulated eQTLs reveal reconfigured interaction landscapes

To overcome the limits of simple linear mediation, we implemented a DNA methylation-modulated eQTL (memo-eQTL) framework [17] that explicitly tests whether proximal CpG methylation, particularly at CpGs within TFBS, modulates the effect of genotype on gene expression (Figure 4A). For each ‘SNP–CpG–gene’ triplet, we fitted three nested models: (M1) a baseline eQTL model (expression ∼ SNP), (M2) an additive model including methylation (expression ∼ SNP + methylation), and (M3) a full interaction model (expression ∼ SNP + methylation + SNP × methylation). We assessed model improvement using LRTs, comparing M3 versus M2 (df = 1) to evaluate the contribution of the SNP × methylation interaction, and M3 versus M1 (df = 2) to assess the combined effect of methylation and interaction terms. *P*-values derived from these tests were corrected for multiple testing using the Benjamini–Hochberg FDR procedure across all tested triplets within each tissue. Triplets were defined as significant memo-eQTLs if both comparisons (M3 vs M2 and M3 vs M1) were significant after FDR correction (FDR < 0.05) (indicated with blue and red dots in Figure 4B).

**Figure 4.**
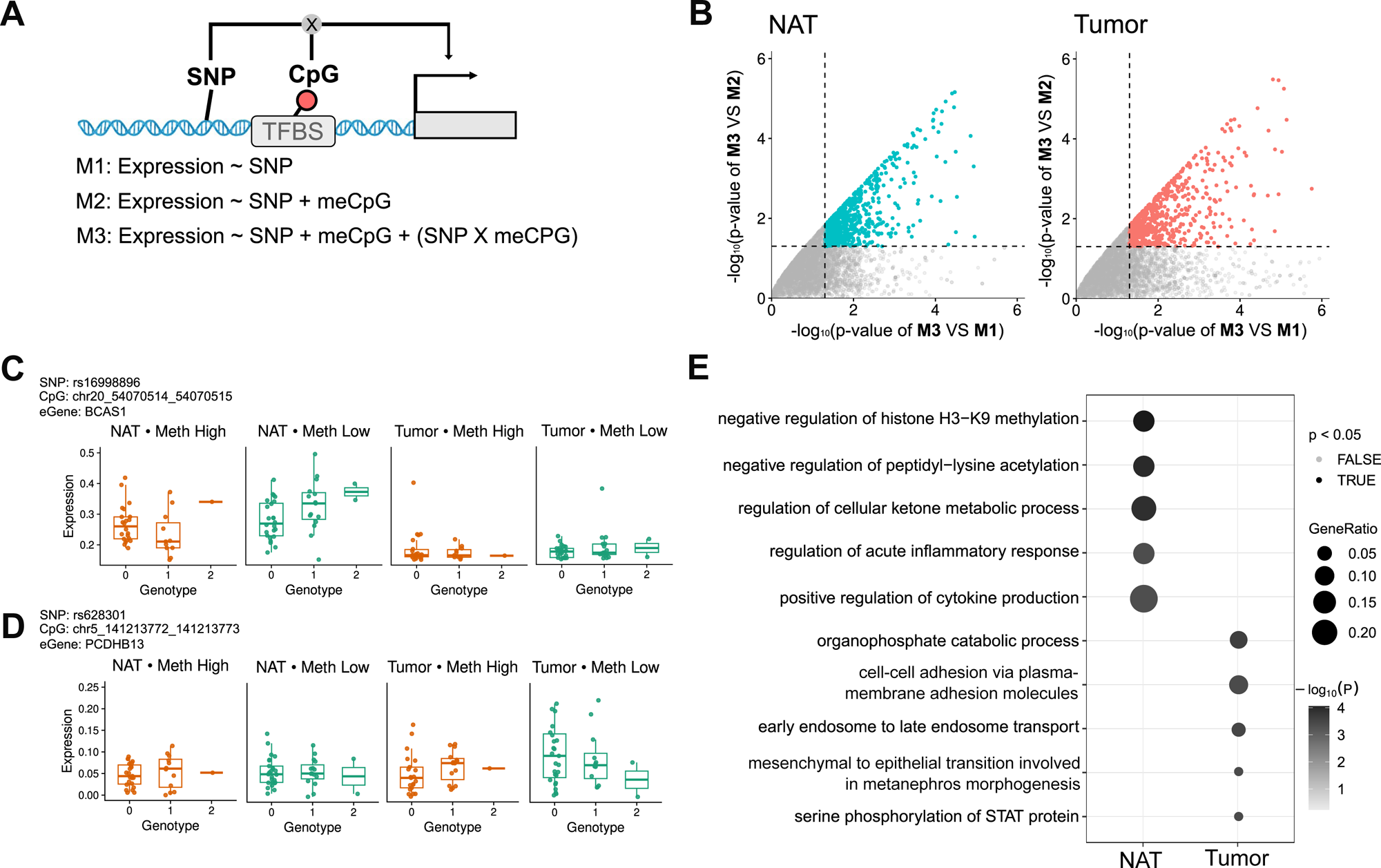
DNA methylation–modulated eQTLs (memo-eQTLs) in normal adjacent and tumor tissues. (**A**) Schematic illustration of the memo-eQTL framework, in which the effect of genotype on gene expression is modeled as a function of proximal DNA methylation through an interaction term. (**B**) Scatter plots summarizing likelihood ratio test results for SNP × methylation interaction effects in NAT (left) and tumor (right) tissues. The x-axis shows −log10(p-value) for the comparison of the full interaction model (M3) versus the baseline model (M1), and the y-axis shows −log10(p-value) for M3 versus the additive model (M2). Each point represents a tested SNP–CpG–gene triplet. (**C, D**) Representative examples of memo-eQTLs identified in NAT (**C**) and tumor (**D**) tissues. Boxplots show genotype-dependent gene expression stratified by high and low methylation states for the corresponding CpG site. Points represent individual samples. (**E**) GO biological process enrichment analysis of memo-eQTL eGenes in NAT and tumor tissues. Dot size indicates gene ratio, and color represents −log₁₀(P-value). Filled dots denote terms with P < 0.05; open dots denote non-significant terms.

We illustrate the architecture of these interactions with two representative loci (Figure 4C–D). For BCAS1 (NAT-specific), the SNP–expression association is evident only under low methylation conditions in NAT and is attenuated at higher methylation levels, with no discernible genotype effect observed in tumor tissue. Conversely, for PCDHB13 (tumor-specific), the SNP shows little or no association with gene expression in NAT, but exhibits a clear genotype-dependent expression pattern in tumor samples, particularly under low methylation conditions. These examples demonstrate that the effect of genotype on gene expression varies depending on local methylation levels and tissue context.

As summarized in Table 1, we applied two FDR thresholds (Q < 0.05 and Q < 0.20) to examine the robustness of memo-eQTL detection. Under both thresholds, tumor tissue consistently harbored more memo-eQTLs than NAT (Tumor/NAT ratio ≈ 4.1 at Q < 0.05 and ≈ 1.8 at Q < 0.20). Notably, the Tumor/NAT ratio narrowed as the threshold was relaxed (from 4.1 to 1.8), suggesting that tumor-specific G×M interactions tend to have stronger statistical signals than their NAT counterparts. Importantly, no memo-eQTL triplets were shared between NAT and tumor tissues, underscoring the complete tissue-specificity of methylation-modulated genetic regulation.

**Table 1.** Summary of promoter-proximal QTL mapping results in NAT and tumor tissues.

| Analysis | N | T | T/N ratio |
| --- | --- | --- | --- |
| eQTL | 944 | 966 | 1.02 |
| mQTL | 643 | 409 | 0.64 |
| **memo-eQTL (Q < 0.05)** |  |  |  |
| eGenes | 18 | 73 | 4.1 |
| Triplets | **222** | **925** | 4.2 |
| Triplets/eGene | 12.3 | 12.7 | 1.03 |
| **memo-eQTL (Q < 0.20)** |  |  |  |
| eGenes | 106 | 190 | 1.8 |
| Triplets | **828** | **1,912** | 2.3 |
| Triplets/eGene | 7.8 | 10.1 | 1.29 |

Triplets:SNP-CpG-gene combinations. N:NAT, T:tumor

A substantial proportion of memo-eQTL eGenes harbored more than 10 significant triplets: 25 of 73 tumor-specific eGenes (34%) and 9 of 18 NAT-specific eGenes (50%), highlighting genes subject to particularly pervasive G×M regulatory control, each influenced by multiple SNP–CpG combinations (Supplementary Table S5 and S6). To characterize the biological functions of these recurrently regulated genes, we performed GO enrichment analysis separately for NAT and tumor eGenes. NAT eGenes were significantly enriched for terms related to epigenetic regulation and immune modulation, including negative regulation of histone H3-K9 methylation, negative regulation of peptidyl-lysine acetylation, regulation of cellular ketone metabolic process, regulation of acute inflammatory response, and positive regulation of cytokine production, consistent with the prominent role of DNA methylation in modulating chromatin state and immune tone in non-malignant tissue. In contrast, tumor eGenes were enriched for distinct biological processes including organophosphate catabolic process, cell-cell adhesion via plasma membrane adhesion molecules, early endosome to late endosome transport, and serine phosphorylation of STAT protein, reflecting processes associated with cancer cell proliferation, invasion, and signaling. The two groups showed largely non-overlapping GO term profiles (Figure 4E), suggesting that methylation-modulated genetic regulation operates through fundamentally different biological pathways in NAT versus tumor tissue.

Having established that memo-eQTLs are tissue-specific, and functionally distinct between NAT and tumor, we next asked whether the magnitude of these G×M interaction effects varies across individual patients and whether such variation carries clinical significance.

### Patient-level SNP-methylation interaction dynamics and clinical significance

To summarize patient-level reprogramming of G×M interactions, we quantified each GxM interaction in each sample (Methods). Figure 5A shows per-sample interaction components (ICs) for all memo-eQTL triplets identified in either NAT or tumor as a heatmap, revealing extensive and largely asymmetric heterogeneity: interactions prominent in NAT are broadly attenuated or absent in paired tumor samples, while a distinct set of interactions emerges de novo in tumors. The low concordance of IC patterns between NAT and tumor tissue across patients confirms that G×M regulatory programs are not merely quantitatively shifted but are qualitatively reconfigured during tumorigenesis.

**Figure 5.**
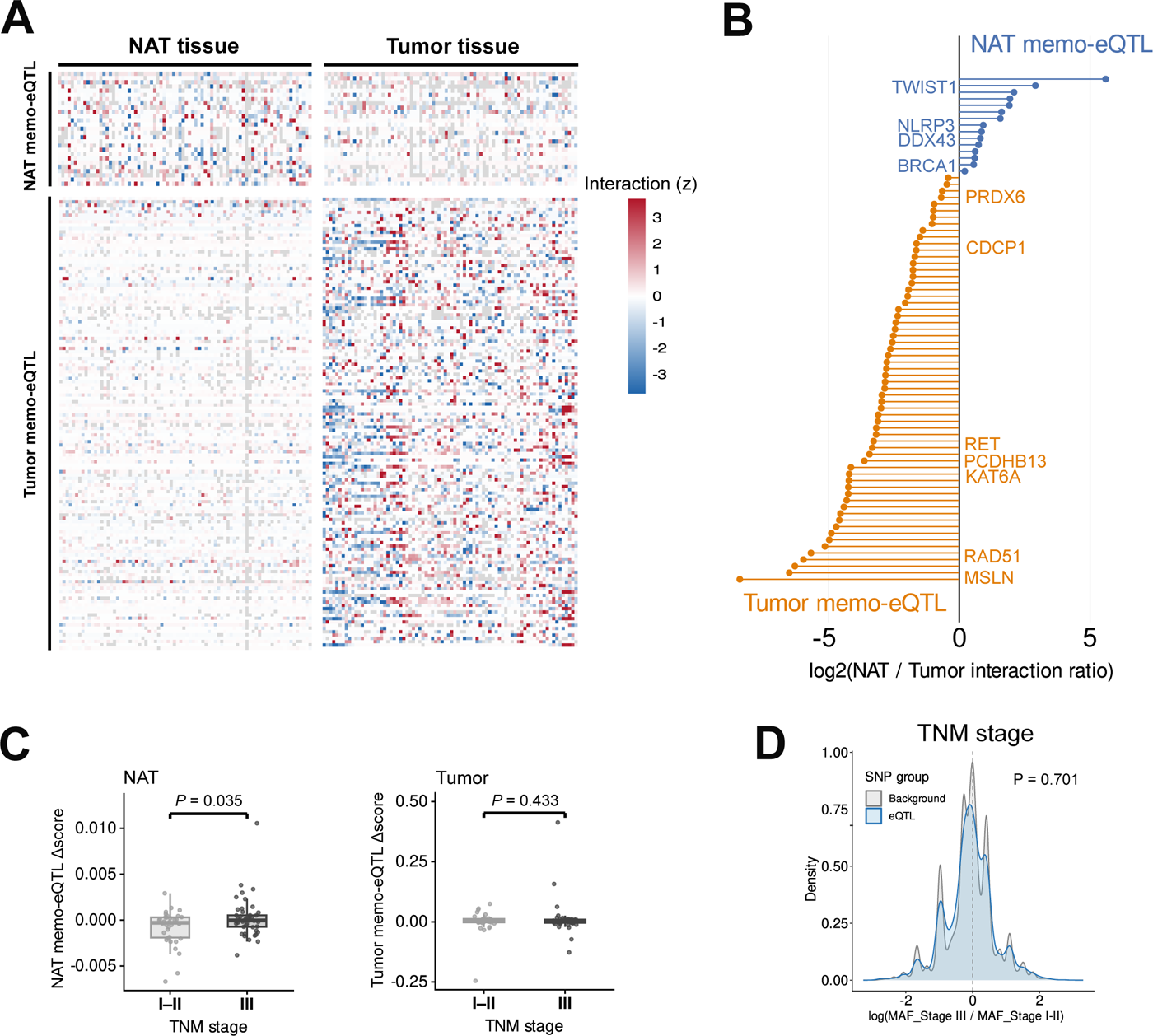
Patient-level heterogeneity of memo-eQTL interaction effects and clinical association. (**A**) Heatmap of patient-specific interaction components (IC) for memo-eQTL triplets. Rows represent individual SNP–CpG–gene triplets, and columns represent patients. Heatmaps are shown separately for NAT-defined and tumor-defined memo-eQTLs. Values are row-wise z-score normalized across patients. (**B**) Gene-level summary of interaction differences between NAT and tumor tissues. Each point represents one memo-eQTL eGene, obtained by aggregating triplet-level interaction components for the corresponding gene. The x-axis shows the log2(NAT/Tumor interaction ratio), where positive values indicate stronger interaction effects in NAT and negative values indicate stronger effects in tumor tissue. Blue points denote NAT-defined memo-eQTL genes and orange points denote tumor-defined memo-eQTL genes. (**C**) Comparison of patient-level memo-eQTL Δscores stratified by TNM stage (I–II vs III). Boxplots show memo-eQTL scores calculated from NAT-defined memo-eQTLs (left) and tumor-defined memo-eQTLs (right). Points represent individual patients. (**D**) Comparison of MAF distributions between eQTL SNPs and background SNPs across clinical subgroups. For each SNP, the log ratio of MAFs was computed between patients stratified by TNM stage (Stage III vs. Stage I–II). Density plots show eQTL SNPs (blue) and background SNPs (grey). *P*-values were determined by two-sided Wilcoxon rank-sum tests.

To quantify the tissue-directionality of these interactions at the gene level, we computed the log_2_ ratio of NAT to tumor IC magnitudes for each memo-eQTL eGene (Methods; Figure 5B). NAT memo-eQTL eGenes, including the EMT regulator TWIST1, the DNA repair gene BRCA1, the inflammasome component NLRP3, and the RNA helicase DDX43, showed consistently stronger interaction effects in NAT than in tumor (positive log_2_ ratios). Conversely, tumor memo-eQTL eGenes, including the mesothelin MSLN, the proto-oncogene RET, the histone acetyltransferase KAT6A, the DNA repair gene RAD51, and the metastasis promoter CDCP1, exhibited markedly stronger interactions in tumor tissue, with several genes showing NAT/Tumor ratios exceeding 10-fold. These robust directional patterns suggest that the same cancer-relevant genes that are subject to methylation-modulated genetic regulation in one tissue context tend to lose that regulation in the other.

We next summarized per-patient interaction burden using a memo-eQTL magnitude Δscore (mean absolute IC across triplets; Methods). Stratifying patients by TNM stage revealed a significant association specifically in NAT: patients with advanced disease (Stage III) having significantly higher NAT memo-eQTL Δscores than those with early-stage disease (Stage I–II) (P = 0.035; Figure 5C, left). By contrast, tumor memo-eQTL Δscores showed no significant stage association (P = 0.433; Figure 5C, right). To exclude the possibility that allele frequency differences between clinical subgroups drive this association, we compared minor allele frequency (MAF) distributions of eQTL SNPs and background SNPs stratified by TNM stage; no significant difference was observed (P = 0.701; Figure 5D), confirming that the clinical signal arises from G×M interactions rather than genotype frequency differences per se. This dissociation is biologically notable: it suggests that the progressive amplification of G×M interactions in histologically normal colonic epithelium, tissue that is not yet transformed, may reflect an epigenetic shift that precedes or accompanies tumor progression. The observation that NAT, rather than tumor tissue, carries a clinically informative G×M interaction signature raises the possibility that methylation-modulated regulatory changes in the tumor microenvironment or adjacent normal tissue could serve as early biomarkers of disease severity.

## Discussion

We investigated how G×M interactions shape gene expression and how these regulatory architectures are reconfigured during the transition from normal to malignant states. By mapping promoter-proximal eQTLs and mQTLs in paired NAT and tumor tissues from CRC patients and performing systematic memo-eQTL analyses, we provide a comprehensive picture of how inherited genetic effects are reorganized during colorectal tumorigenesis. Leveraging paired samples to control for germline background, we show that genetic regulation is not simply lost in cancer; rather, it is fundamentally reorganized through complex, context-dependent G×M interactions.

Traditional regulatory models often assume a linear cascade in which DNA methylation mediates genotype effects on expression. However, our mediation analyses (Figure 3) demonstrate that the canonical ‘SNP → methylation → expression’ pathway is uncommon in both normal and malignant colorectal tissues, revealing a substantial mediation gap. To capture the complexity missed by simple mediation, we applied a memo-eQTL framework that explicitly models G×M interactions [17]. As shown in Figure 4B, the full interaction model (M3), which includes the SNP × methylation term, consistently outperformed the reduced additive models (M1 and M2) for most SNPs, indicating that many genetic effects are conditional on the local methylation state.

A notable outcome of this reorganization is its functional specificity. Most G×M interactions arise de novo in tumors, where interaction magnitudes (i.e., ICs) tend to be larger. Although the greater expression and methylation variability in tumors may partly contribute to increased detection power, this does not diminish the biological relevance of the observed interactions. This suggests that the somatic epigenetic landscape can unlock latent germline influences on pathways that promote malignant phenotypes, effectively creating a tumor-specific regulatory niche in which inherited variants acquire new functional consequences.

One striking finding is that the intensity of interaction-based regulation in the NAT, rather than the tumor itself, better reflects clinical progression (Figure 5). Patients with Stage III disease exhibited significantly higher NAT memo-eQTL Δscores than those with Stage I–II (*P* = 0.035), consistent with our previous work showing that NAT transcriptomes are more informative than tumor transcriptomes for predicting CRC recurrence [16]. These findings align with the field-cancerization concept: histologically normal tissue adjacent to tumors can undergo molecular priming and regulatory remodeling that mirror tumor aggressiveness [35]. Because tumors are often highly heterogeneous and noisy, disrupted genotype–methylation dynamics may be easier to detect in the relatively more stable NAT, offering a clearer readout of the patient’s clinical state.

Our findings have significant implications for interpreting genetic association signals in cancer. First, because regulatory effects are highly context dependent, a variant’s effect size and direction may change across tissue states. Functional annotation of GWAS loci in cancer should therefore incorporate local epigenetic context and state-specific interactions. Second, the memo-eQTL score raises the possibility that normal-tissue biopsies could carry prognostic information by reporting the extent of epigenetic–genetic dysregulation.

Despite these insights, several limitations remain. It is currently unclear whether disrupted G×M interactions drive progression or are consequences of tumor-associated microenvironmental changes; temporal or perturbational studies will be required to establish directionality. Furthermore, as our analyses relied on bulk RNA-seq and bisulfite-seq data, we cannot assign reconfigured interactions to malignant epithelial cells versus stromal or immune compartments; single-cell or spatial multi-omics will be essential to resolve cellular origin. Local somatic events, such as copy-number changes or loss of heterozygosity, may also alter genotype calls or regulatory relationships. A further limitation concerns statistical power. Split-half reproducibility analysis revealed that within-tissue eQTL overlap at N = 40 was markedly low (Jaccard ∼0.02; Supplementary Figure S1A), indicating that individual eQTL identification is sensitive to sample composition at this cohort scale. While aggregate-level metrics, including the saturation curve and per-eQTL reproducibility fractions, support the overall validity of our eQTL calls (Figure 1D, E), the incomplete detection of individual regulatory associations is an inherent consequence of the modest sample size, and larger cohorts will be needed for more comprehensive eQTL characterization. Future studies integrating comprehensive somatic profiles and larger, well-annotated cohorts will be necessary to robustly link these interaction patterns to clinical outcomes across diverse cancer types.

## Conclusions

In summary, our paired NAT–tumor memo-eQTL analysis shows that cancer rewires genotype-driven regulation through context-dependent G×M interactions. These results underscore the need to model regulatory interactions, beyond marginal effects or simple mediation, to accurately map genetic influences and their clinical consequences in cancer.

## Declarations

### Ethics approval and consent to participate

Not applicable

### Consent for publication

Not applicable

### Availability of data and materials

The summary statistics of eQTL, mQTL, and memo-eQTL analyses generated in this study are provided as Supplementary Tables (Additional Files 1-7).

### Authors’ contributions

Byounghun Kim: Formal analysis; Investigation; Data curation; Visualization; Writing – original draft.

Hanbeen Kim: Methodology; Data curation; Formal analysis; Software; Writing – review & editing.

Min-Kyeong Kwon: Formal analysis; Software; Validation; Writing – review & editing.

Sridhar Hannenhalli: Supervision; Writing – review & editing.

Sun Shim Choi: Conceptualization; Supervision; Funding acquisition; Writing – review & editing; Project administration.

## Acknowledgements

Not applicable

## Additional files

**Additional file 1:**

Supplement Table S1. Promoter-proximal eQTLs identified in NAT and tumor tissues of colorectal cancer patients.

Description: Significant SNP–gene pairs from promoter-proximal eQTL analysis, including gene annotations, SNP genomic coordinates, and promoter boundary definitions (NAT-specific, tumor-specific, or shared).

**Additional file 2:**

Supplementary Table S2. Promoter-proximal mQTLs identified in NAT and tumor tissues of colorectal cancer patients.

Description: Significant SNP–gene pairs from promoter-proximal mQTL analysis, including CpG annotations, SNP genomic coordinates, and promoter boundary definitions (NAT-specific, tumor-specific, or shared).

**Additional file 3:**

Supplementary Table S3. Pathway enrichment analysis of tissue-specific and shared proximal eQTL-associated genes.

Description: Pathway enrichment results for eGenes classified as NAT-specific, tumor-specific, or shared. Results include pathway identifiers, gene ratios, and p-values from KEGG over-representation analysis.

**Additional file 4:**

Supplementary Table S4. Pathway enrichment analysis of tissue-specific and shared proximal mQTL-associated genes.

Description: Pathway enrichment results for eGenes classified as NAT-specific, tumor-specific, or shared. Results include pathway identifiers, gene ratios, and p-values from KEGG over-representation analysis.

**Additional file 5:**

Supplementary Table S5. memo-eQTLs identified in NAT and tumor tissues at stringent and lenient significance thresholds.

Description: Significant SNP–CpG–gene triplets from memo-eQTL analysis in NAT and tumor tissues, identified at two FDR thresholds (Q < 0.05 and Q < 0.2).

**Additional file 6:**

Supplementary Table S6. Gene-level summary of memo-eQTL triplet counts in NAT and tumor tissues.

Description: Gene-level aggregation of significant memo-eQTL triplets, ranked by the number of SNP–CpG–gene triplets per gene in descending order.

**Additional file 7:**

Supplementary Table S7. Gene Ontology enrichment analysis of memo-eQTL-associated genes in NAT and tumor tissues.

Description: Gene Ontology over-representation analysis results for memo-eQTL eGenes identified at FDR < 0.05 in NAT and tumor tissues.

**Additional file 8:**

Supplementary Figure S1. Split-half eQTL reproducibility and eSNP allele frequency analysis. Description: Within-tissue split-half Jaccard index distributions (40 vs. 40 patients, 100 iterations)

