## Supplementary figure 1 for "Genotype and methylation interact to reconfigure transcriptional regulation in colorectal cancer"

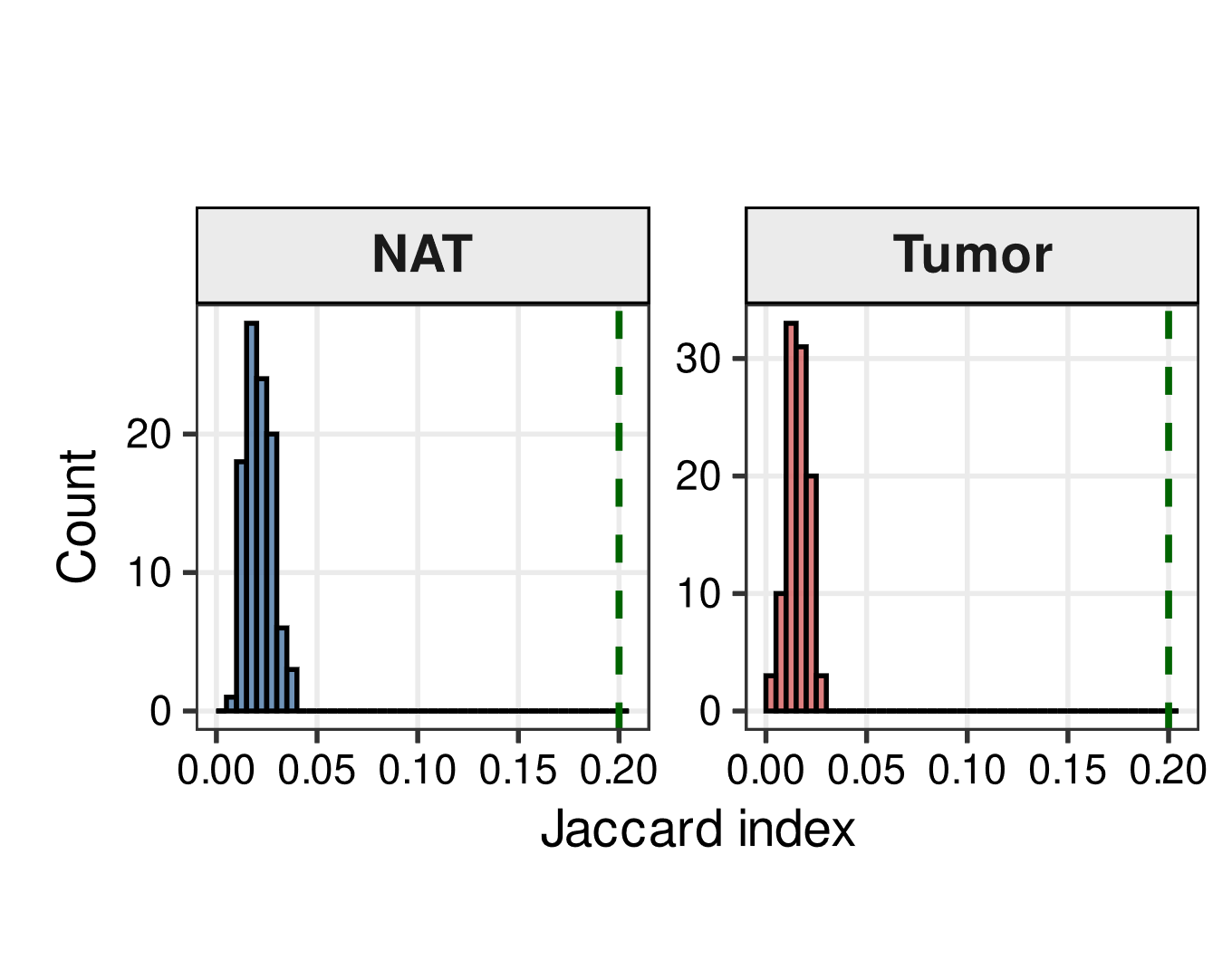

**Supplementary Figure S1.** **Split-half eQTL reproducibility and eSNP allele frequency analysis.**

(**A**) Distribution of within-tissue split-half Jaccard indices across 100 iterations. In each iteration, 80 patients were randomly divided into two mutually exclusive groups of 40, and promoter-proximal eQTL mapping was performed independently for each group within the same tissue. The Jaccard index between the two resulting eQTL sets was computed for NAT (left) and Tumor (right). Green dashed lines indicate the between-tissue Jaccard index (~0.20) obtained from the full cohort (N = 80; Figure 1G) for comparison.
